# Population genomic analysis reveals genetic structure and thermal-tolerant genotypes in remnant Tasmanian giant kelp populations

**DOI:** 10.1101/2023.10.08.561451

**Authors:** Cintia Iha, Cayne Layton, Carlos E. Amancio, Warren Flentje, Andrew Lenton, Ceridwen I. Fraser, Craig Johnson, Anusuya Willis

**Author notes:** **Correspondence:** Cintia Iha. Text formatted in British English.

## Abstract

Giant kelp, *Macrocystis pyrifera*, is a foundation species that forms dense forests of complex physical habitat and supports coastal biodiversity, productivity, and other essential ecosystem services. Tasmanian coasts have suffered a massive decline in giant kelp forests due to changes in regional oceanography and environmental conditions, but efforts are being made to restore these disappearing populations using identification and selective breeding for lineages that are more tolerant to warmer temperatures. Here, we used gametophytes that originated from remnant populations collected at three sites in northeastern and three sites in southeastern Tasmania to determine the genetic structure of the giant kelp population via genotyping-by-sequencing (GBS) and assembled a draft genome from a Tasmanian giant kelp sporophyte individual. Previous research identified seven strain lines that were tolerant to warm temperatures, and we used the GBS data to test whether genotypes were associated with warm tolerance. Kelps from the north were genetically distinct from the southern ones, with much lower gene flow between regions than within regions. These results revealed that giant kelp populations from Tasmania are more genetically structured than previously thought. Two loci were significantly associated with warm temperature tolerance. They are population-specific: the alternative allele from one of the loci was found only in the northern populations, while the other was found in the southern populations. This could suggest that thermal tolerance is adapting locally or drifting given large changes in population demography, but further research is needed to confirm this hypothesis. Our research sheds light on genetic patterns in this critical habitat-forming kelp and will help inform conservation management, including selective breeding.

## 1 Introduction

Giant kelp, *Macrocystis pyrifera* (Laminariales, Ochrophyta), is a foundation species that can rapidly grow to over 40 m tall and forms dense kelp forests with floating surface canopies. These forests create complex habitats that support coastal productivity and provide essential ecosystem and marine-related economic services (Bennett *et al*., 2016). It is found along the North American west coast and across temperate and sub-Antarctic coasts in the Southern Hemisphere (Schiel & Foster, 2015). The giant kelp struggles to grow in temperatures above 20°C and low nutrient environment (Schiel & Foster, 2015). Kelp forests are declining in many parts of the world, except perhaps in some polar and sub-polar regions (Krumhansl *et al*., 2016, Mora-Soto *et al*., 2021), due to changes in regional oceanography and environmental conditions, including warming waters and reduced coastal nutrients.

Tasmanian shallow-water ecosystems were once home to dense giant kelp forests, but they have declined dramatically, with at least 95% lost in recent decades (Johnson *et al*., 2011, Steneck & Johnson, 2014, Butler *et al*., 2020). The disappearance of these kelp forests is associated with an increase of at least ~2°C in the sea surface temperature (SST) and low nutrient concentrations in surface water in Tasmania (Johnson *et al*., 2011, Butler *et al*., 2020). The water temperature of the northern region is warmer, with the average SST ranging between 12°C to 15.5°C, while the SST at ~200 km to the south is 1-2°C colder. The Tasmanian east coast is also a global warming hotspot for marine heatwaves due to the influence of the East Australian Current (EAC), which is a warm and nutrient-depleted current (Oliver & Holbrook, 2014, Butler *et al*., 2020). Giant kelp populations from the northern region have been suffering more from the effects of warmer temperatures than southern sites. Efforts are now being made to restore these disappearing kelp forests by seeding populations with material from remnant locations and selective breeding programmes (Layton *et al*., 2020, Layton & Johnson, 2021). Reproductive tissues from mature sporophytes have been collected from giant kelp populations in the northern and southern regions of the eastern coast of Tasmania. A gametophyte seed bank has been established from those giant kelp reproductive tissues, and these gametophytes have been used for reforestation experiments (Layton & Johnson, 2021). Reforestation attempts have demonstrated that the viability of sporophytes decreases significantly with increasing water temperatures but that some strain family lines present an increased tolerance to warmer waters. These studies have identified individuals within populations that can survive in temperatures 4°C above their normal range and have been the focus of the reforestation efforts (Layton & Johnson, 2021).

Despite the importance and previous abundance of giant kelp on the Tasmanian coast, little is known about the genetic structure of the remnant populations, particularly the potential warm-tolerant individuals. Most of the existing studies on the population genetics of giant kelp from the Southern Hemisphere are based on a few molecular markers or microsatellites (Coyer *et al*., 2001, Macaya & Zuccarello, 2010, Astorga *et al*., 2012, Durrant *et al*., 2015, Camus *et al*., 2018). The most common phylogeographic markers, chloroplast *rbc*L and mitochondrial *cox*1 show little variation for *Macrocystis* in southern Australia (Durrant *et al*., 2015). Genetic studies using nuclear microsatellite loci from northeastern or southeastern Pacific populations showed higher genetic variability even among close sites (Alberto *et al*., 2010, Johansson *et al*., 2015, Camus *et al*., 2018). However, information from such small genome regions is insufficient to allow inferences linking organism adaptation to specific phenotypes (e.g., tolerance to warm temperatures). Genotyping-by-sequencing (GBS), which uses restriction enzymes to isolate random fragments of genomic DNA, is a cost-effective method that reveals thousands of genome-wide polymorphisms, potentially enabling the discovery of loci associated with phenotypic traits (Narum *et al*., 2013). Scanning genetic markers across the whole genome, as commonly performed with genome-wide association studies (GWAS), is useful for investigating genetic signals of tolerance to warm waters or any other phenotype that might help the conservation of species. Conservation genomics is a field that uses genomic approaches to assist conservation efforts (Ouborg *et al*., 2010), such as the reintroduction of selective populations (He *et al*., 2016). GBS could, therefore, detect genetic structure in giant kelps and could also reveal genotypes associated with warm-tolerant individuals.

Given the major and rapid decline of giant kelp forests across Tasmania and future forecasts for ongoing warming—including marine heatwave events (Li *et al*., 2020)—tolerance to warming waters will likely be essential for this important species and habitat type to survive in Australian waters. Understanding the genetic structure of the remnant populations and the genetic basis for thermal tolerance is crucial for building a management plan for giant kelp forest conservation and reforestation. This study was thus performed aiming to determine the genetic structure of remnant giant kelps from Tasmania via genotyping-by-sequencing (GBS) and to test whether any genotypes are potentially associated with warm tolerance. Because of its high power for phylogeographic research, including in other marine macroalgae (Fraser *et al*., 2016), we hypothesised that GBS would detect hitherto unknown spatial genetic structures among populations in Tasmania and that some loci would be linked to warm tolerance.

## 2 Material and Methods

### 2.1 Nuclear Genome Sequencing

Using a reference genome for SNP calling can result in higher quantity SNPs and ensure the SNPs are not from potential contaminants. During the development of this study, there was no giant kelp genome available for the southwestern Pacific, so we sequenced and assembled the draft nuclear genome of a Tasmanian *M. pyrifera* sporophyte.

#### 2.1.1 Collection of the sporophyte, DNA extraction and sequencing

We collected ~30 cm of *Macrocystis pyrifera* sporophyte apical lamina from Blackmans Bay beach, Tasmania, Australia (43°01’02.4”S, 147°19’45.8”E). The tissue was rinsed in deionised water and dried on absorbent paper. Small pieces of ~3 cm² were then cut from the thinnest portions of the lamina and stored in silica gel. Qiagen DNeasy PowerPlant Pro Kit was used for DNA extraction, following the modified method outlined by Peters *et al*. (2020). Samples were soaked for 24 h at 65°C in 500 µl of PowerBead solution and 3 µl of RNase A (10 mM). After incubation, 100 µl of 100% isopropanol was added and incubated at 65°C for 90 mins, with vortexing every 30 mins. Samples were then placed in a FASTPREP-24™ 5G homogeniser (MP Biomedicals, Santa Ana, California, USA) for 40 seconds. Subsequent steps followed the manufacturer’s protocols, with elution of DNA in 100 µl of TE buffer, incubated for 10 min before centrifugation. We measured the DNA quantity with QUBIT with dsDNA Broad Range (BR) kit and quality on Nanodrop.

The DNA library preparation and whole-genome sequencing were carried out by the Australian Genome Research Facility (AGRF; Melbourne, Australia). Sequencing was performed on an Illumina NovaSeq 6000 using PE 150 bp High Output kit (Illumina, San Diego, California, USA).

#### 2.1.2 Genome and transcriptomes assembly

The nuclear genome was assembled with MaSuRCA v4.0.5 (Zimin *et al*., 2013) using raw reads with SOAP assembly mode generating 4713 scaffolds. The scaffolds were mapped with Bowtie 2 v.2.3.4 (Langmead & Salzberg, 2012) to check the depth coverage and coverage information for each scaffold that was obtained with pileup.sh script from BBtools v.38.37 (Bushnell, 2021). BUSCO v5.1.2 (Waterhouse *et al*., 2018) was used to check genome completeness.

To assist the genome annotation and remove contaminant scaffolds, we used two transcriptomes from *Macrocystis pyrifera* available on GenBank: PRJNA322132 (Salavarria *et al*., 2018) and PRJNA353611. The transcriptomes were assembled individually with Trinity v2.12.0 (Grabherr *et al*., 2011) using both *de novo* and genome-guided modes. *De novo* mode assemblies were used to guide the identification of contaminant sequences described below. The genome-guide mode was assembled using partially cleaned genome scaffolds generated in this study.

#### 2.1.3 Removal of contaminant sequences

We performed a comprehensive strategy to identify sequences of putative contaminants that combine GC content, read coverage, taxonomical annotation, and transcriptome data. Blobtools v1.1.1 (Laetsch & Blaxter, 2017) was employed to integrate the taxonomic identification, GC content and read coverage for each scaffold. The taxonomic identification was performed by comparing the scaffolds with Uniprot, Silva and a custom database. The custom database was created by retrieving protein sequences of bacteria, archaea, fungi, viruses, Stramenopiles, Rhodophyta and Chlorophyta from the GenBank non-redundant database. We used Diamond v.2.0.4 blastx (Buchfink *et al*., 2015) to align the sequences from the custom and Uniprot database against the genome scaffolds, using ten max target sequences, sensitive mode, and e-value of 1e^-25^. We used BLASTn v.2.12.0 to align the Silva database against genome scaffolds using an e-value of 1e^-65^.

From the scaffolds with taxonomic designation assigned as Eukaryote, 3339 matched with *Ectocarpus* (a brown algae genus), and 19 had other taxa. We manually inspected the non-*Ectocarpus* sequences before removing them as putative contaminations. For the scaffolds with taxonomic designation assigned as Bacteria or “no-hit”, we used *Macrocystis pyrifera* transcripts assembled with Trinity using the *de novo* method to map against the scaffolds. Scaffolds with mapped transcripts and indications of introns were considered as *M. pyrifera* scaffolds and retained. Scaffolds with no indication of introns were removed as putative contaminations, irrespective of whether they had mapped transcripts or not. For the scaffolds without mapped transcripts, we removed the ones that are GC content and read coverage outliers (above and below the upper and lower quartile, respectively).

#### 2.1.4 *Ab initio* prediction of protein-coding genes

We adapted the workflow for *ab initio* prediction of protein-coding genes from Chen *et al*. (2020). Repetitive elements in the genome assembly were first predicted with *de novo* mode using Repeat Modeler v2.0.2a (http://www.repeatmasker.org/RepeatModeler/). These repeats were combined with known repeats in the RepeatMasker database (release 20201109) to generate a custom repeat library. All repetitive elements in the assembled genome scaffold were then masked using RepeatMasker v4.1.2 (http://www.repeatmasker.org/) using the custom repeat library, creating two scaffold data with hard-masked repeats and soft-masked repeats.

Vector sequences were removed from the assembled transcripts using SeqClean (Chen *et al*., 2007) based on the UniVec (build 10) database. The PASA pipeline v2.5.1 (Haas *et al*., 2003) was used to predict protein-coding genes using unmasked repeat scaffolds and the vector-trimmed transcriptome assemblies, allowing a max intron length of 70000. The predicted protein sequences from multi-exon transcript-based genes with complete 5’ and 3’-ends were searched with BLASTp (evalue 1e^-20^) against the non-redundant database. Genes with significant BLASTp hits (>80% query coverage) were retained. Transposable elements were identified with Transposon-PSI (http://transposonpsi.sourceforge.net/).

Proteins putatively identified as transposable elements were removed. Those remaining were clustered using CD-HIT v4.8.1 (Li & Godzik, 2006) with ID=75% to yield a non-redundant protein set, and the associated transcript-based genes were kept. These genes were further processed by the Prepare_golden_genes_for_predictor.pl script from the JAMg package (https://github.com/genomecuration/JAMg). This step yielded a set of high-quality “golden” genes, which were used in SNAP (Korf, 2004) with soft-masked scaffolds at the default setting. Soft-masked scaffolds were also used for AUGUSTUS (Stanke *et al*., 2006) with prediction mode. We employed GeneMark-ES v4.30 (Lomsadze *et al*., 2018) at default settings to generate predictions from the hard-masked scaffolds. Subsequently, all genes predicted using GeneMark-ES, PASA, SNAP and AUGUSTUS were integrated into a combined set using EvidenceModeler v1.1.1 (Haas *et al*., 2008), following a weighting scheme of GeneMark-ES 2, SNAP 2, Augustus 6 and PASA 10.

### 2.2 Genotyping-by-sequencing

#### 2.2.1 Macrocystis pyrifera gametophytes

We obtained gametophyte cultures from the long-term seed bank of *M. pyrifera* stored at the Institute for Marine and Antarctic Studies, University of Tasmania (Layton & Johnson, 2021). These gametophyte cultures were released from sporophyte individuals, containing both male and female gametophyte cells, and kept at 4°C under red light to prevent sexual reproduction. We obtained six to eight cultures (herein referred to as ‘individuals’ because the genotype of each culture represents its parental sporophyte) from three geographic population sites collected on the northeast coast (A, B, and C) of Tasmania and three on the southeast coast (D, E and F), totalling six populations and 45 individuals (Fig. 1).

**Fig. 1.**
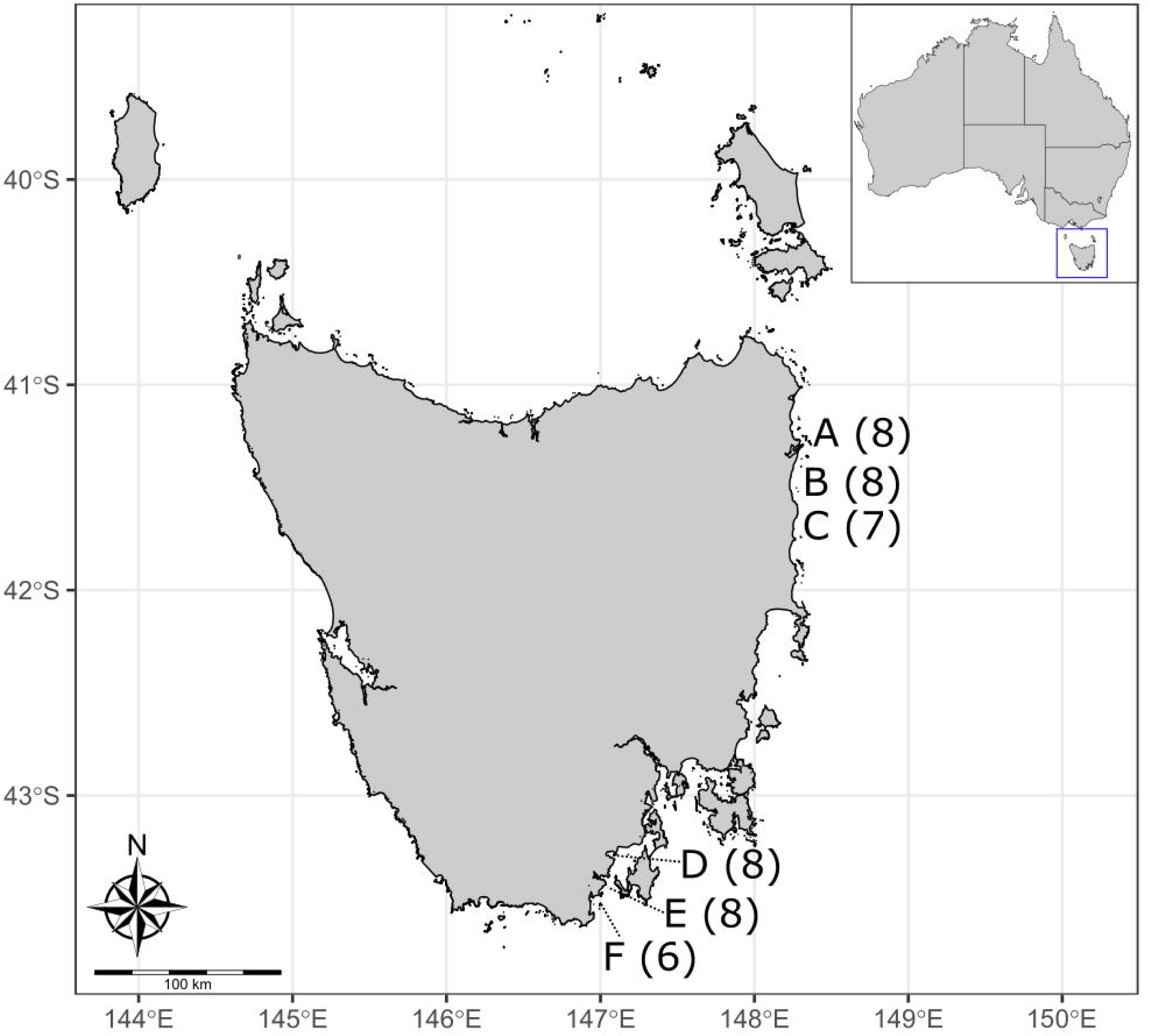
Map of Tasmania showing the collection spots. The number of individuals is indicated in parentheses. (Adapted from Layton & Johnson, 2021)

#### 2.2.2 DNA extraction and ddRAD-seq sequencing

We filtered ~120-200 mg of fresh gametophyte cells using PC filter papers that were scraped with a stainless-steel spatula previously washed and cleaned with 70% ethanol to avoid contamination. The DNA extraction, quality and quantity checks were performed following the same protocol for sporophytes described above. Genotyping-by-sequencing library preparation was performed by AGRF, with a ddRAD-based protocol (Peterson *et al*., 2012) for processing the samples. The restriction enzymes were PstI and MseI. Sequencing was performed on an Illumina NovaSeq 6000 with a 150bp single-end run.

#### 2.2.3 SNP calling and population genetic analysis

Raw reads were demultiplexed with Stacks v2.3d (Catchen *et al*., 2011). All samples were quality-checked with FastQC v0.11.9 (Andrews, 2010) and visualised with MultiQC v1.11 (Ewels *et al*., 2016). We used the reference-map pipeline from Stacks to identify loci and estimate genotypic variation. We first mapped the GBS reads of each sample against the reference genome with BWA v0.7.17 (Li & Durbin, 2009), then performed the SNP calling using the pipeline of *gstacks* component from Stacks with default settings using a population map indicating the population (A, B, C, D, E, and F) and region (north or south) for each individual to identify loci. We used the *populations* component of Stacks to filter SNPs with a minimum threshold of at least 65% of samples (-r 0.65). SNP data were again filtered using vcftools v0.1.16 (Danecek *et al*., 2011), setting minor allele frequency to greater than 0.02, minimum missing data to 0.6, minimum of 5 and maximum 250 mean depth for a site and depth allowed for a genotype. The *populations* component of Stacks was used to estimate the mean nucleotide diversity (π), mean expected heterozygosity (*H_E_*), and mean observed heterozygosity (*H_O_*) within each sampling site using the filtered SNP data.

All other statistical analyses were conducted in R (R Core Team, 2018). We used POPPR (Kamvar, Tabima *et al.,* 2014) to calculate a hierarchical molecular variance analysis (AMOVA) using Region → Population hierarchy. Pairwise *F_ST_* between populations was calculated with STAMPP (Pembleton *et al*., 2013) with 10,000 permutations. To investigate population structure, we performed a principal component analysis on the filtered SNPs with the glPCA() function from adegenet package v2.1.5 (Jombart & Ahmed, 2011). First, we used the find.cluster() function to determine the number of groups (K) that were estimated in our samples, adding the “diffNgroup” criterion and repeating the analysis ten times. The optimal K was selected based on automated detection. To determine whether the geographic collection location reflected the genetic populations, we performed a Discriminant Analysis of Principal Components (DAPC) with an assignment based on geographic sampling location. We further explored population structure by undertaking DAPC analysis (Jombart *et al*., 2010). DAPC analysis was performed with two principal components (n.pca = 2) and two discriminant analysis eigenvalues (n.da = 2). The DAPC posterior probability values, defining membership probability to population individuals, were plotted.

#### 2.2.4 *De novo* assembly and model for SNP association with tolerance to warm temperatures

Seven strain lineages in our sampling presented some tolerance to warm temperatures in previous study: B7, C2, C3, C8 from the northern region and D2, E5, and E6 from the southern region (Layton & Johnson, 2021). The SNP calling based on alignment to the reference genome resulted in few SNPs shared among all 45 individuals. Using the same methodology described below, no SNPs associated with warm temperature tolerance were found (data not shown). Therefore, we performed another approach to identify more SNPs.

We performed a *de novo* assembly with Stacks, indicating the region (north or south) to map each individual. Eleven individuals were excluded due to low-depth read coverage, and the new dataset comprised 34 individuals (including the seven individuals from thermal tolerant strain lines), of which 15 were from the northern region and 19 from the south. We used the *gstacks* component of Stacks for SNP calling with default parameters. The SNPs were filtered using vcftools excluding sites with more than two missing genotypes over all individuals and with Phred quality below 30 and a minimum of 5 mean depth.

To identify possible SNPs associated with tolerance to warm temperatures, we used GEMMA v0.98.5 (Zhou & Stephens, 2012) to perform a univariate linear mixed model analysis to test marker association for a single binary phenotype (tolerant versus non-tolerant) to account for population structure (north and south) using filtered *de novo* SNP dataset of 34 individuals. A relatedness matrix was required for the analysis. We used a standardised matrix (-gk 2) to look for potential SNPs with lower minor allele frequency (MAF). The univariate linear mixed model performing the likelihood ratio test was conducted with “-notsnp” flag that disables the minor allele frequency cutoff. As the genetic structure was strong in these populations, we included a covariate file indicating the genetic group of the individuals, north or south region. We used qvalue v.2.30.0 (Storey *et al.,* 2022) to adjust the likelihood ratio test p-values with a false discovery rate (FDR) estimation.

## 3 Results

### 3.1 Draft nuclear genome

The draft nuclear genome for *M. pyrifera* had 3,864 scaffolds after removing putative contaminant sequences. The total length of assembled bases was 52 MB, N50 length of 26,628 KB, an average depth coverage of 1064.34⨉ and GC content of 50.31%. Gene annotation resulted in 2,950 predicted protein-coding genes. The completeness of the genome was checked using BUSCO analysis, resulting in 12.9% of completed BUSCOs for the Eukaryota database and 11% for the Stramenopiles database. Assembling a high-quality nuclear genome was beyond the scope of this study, but despite the low genome completeness, it was helpful for improving SNP calling and removing contaminant reads. The draft genome and predicted proteins are deposited at https://figshare.com/s/10d87c46afe8d98f8eea (*This is a temporary link for the review process. A public DOI will be generated with the acceptance of the manuscript*).

### 3.2 Genetic structure

Two samples from population A were removed prior to analysis due to low coverage. The SNP-calling, based on alignment to the reference genome, generated 21,051 variant loci located in 2,232 scaffolds. The average of polymorphic loci across in all sites was 4.6%. After filtering, we obtained 5,911 SNPs located in 1,505 scaffolds. Observed heterozygosity (*H_O_*) was higher than expected heterozygosity (*H_E_*) for all populations (Table 1), with *H_O_* ranging from 0.18 to 0.20 compared to *H_E_* 0.12 to 0.15. Nucleotide diversity was low and similar among all populations. Hierarchical AMOVA found strong and significant genetic differentiation (ϕ_ST_ = 0.65, p = 0.01), with 55% of the variance was explained between north and south regions (Table 2). The pairwise F_ST_ between populations within a region was in general low, ranging from 0.029 to 0.122, with the highest value between southern sites D and F, and the lowest value between northern sites B and C (Fig. 2). Populations between regions (north or south) presented higher pairwise F_ST_, ranging from 0.255 to 0.31 (Fig. 2). All comparisons were significant (p < 0.05).

**Fig. 2.**
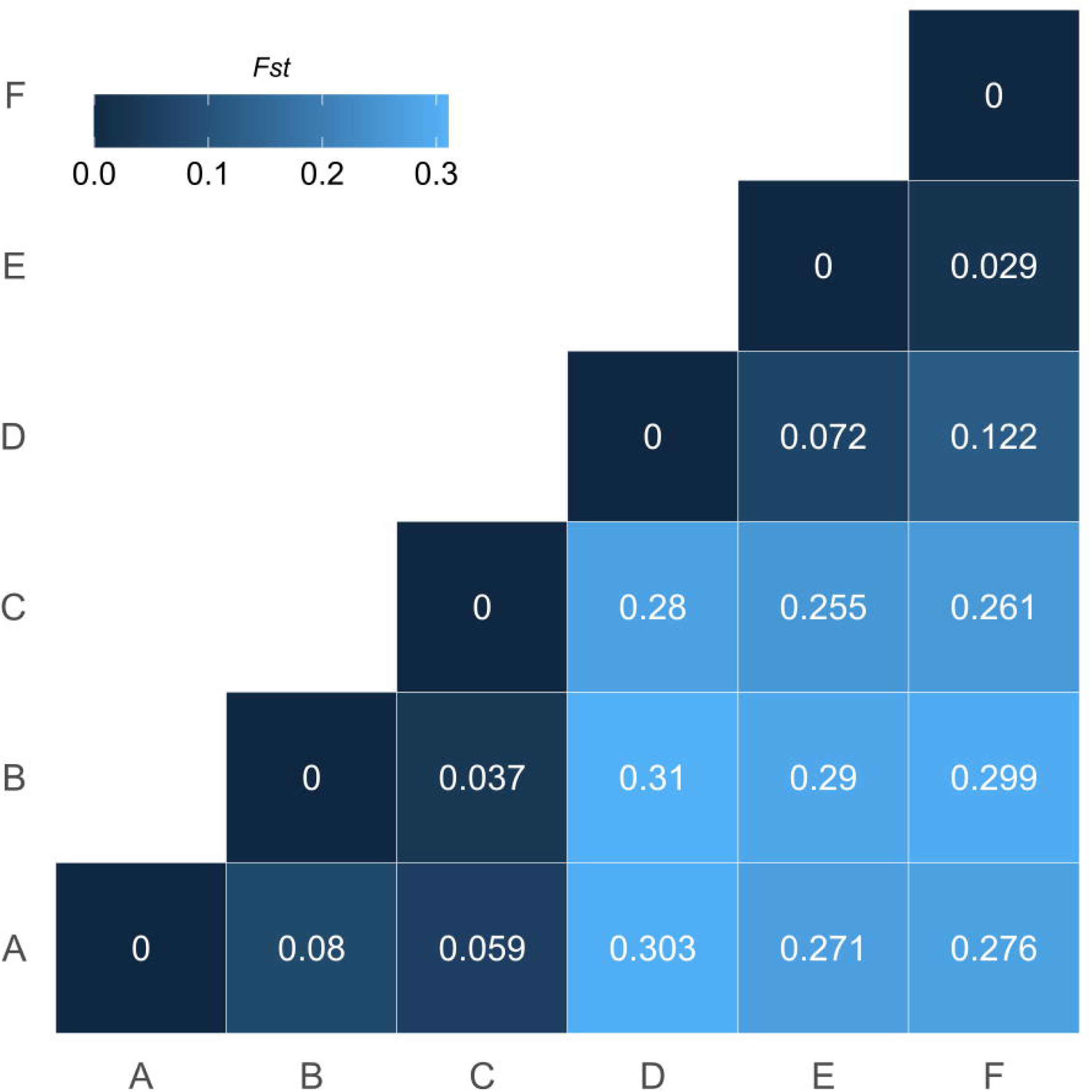
Pairwise F_ST_ estimates between all populations.

**Table 1 –.** Genetic metrics for giant kelp populations on the eastern coast of Tasmania for filtered dataset (5,911 SNPs).

| Region | Population | $H_O$ | $H_E$ | Nucleotide diversity ($\pi$) |
| --- | --- | --- | --- | --- |
| North | A | 0.19 | 0.13 | 0.14 |
| North | B | 0.18 | 0.12 | 0.13 |
| North | C | 0.20 | 0.14 | 0.14 |
| South | D | 0.18 | 0.12 | 0.13 |
| South | E | 0.18 | 0.13 | 0.13 |
| South | F | 0.20 | 0.15 | 0.16 |

**Table 2 -.** Hierarchical analysis of molecular variance (AMOVA) of filtered SNP data set.

| | Degrees of Freedom | Sum of Squares | Mean of Squares | Variance ($\sigma$) | % Total Variance | p-value |
| --- | --- | --- | --- | --- | --- | --- |
| Between Region | 1 | 2196.29 | 2196.29 | 93.43 | 54.60 | 0.01 |
| Between population Within Region | 4 | 743.31 | 185.83 | 17.69 | 10.34 | 0.01 |
| Within population | 37 | 2220.44 | 60.01 | 60.01 | 35.07 | 0.01 |
| Total | 42 | 5160.03 | 122.86 | 171.13 | 100.00 |  |

The PCA showed strong differentiation between populations from the north and south of Tasmania, with PC1 explaining 32.7% of variation (Fig. S1) with less variance within regions (PC2 = 6.2%). Northern populations (A, B and C) are more genetically similar to each other than the southern populations, where population D was more genetically distant from E and F. Although K=3 was also somewhat supported, the sharp decrease of the Bayesian Information Criterion (BIC) values from K=1 to K=2 from the k-means clustering (Fig. 3A) indicates the existence of two major genetic clusters in this study. The distribution of the group membership (*grp*) for each sample indicates two clusters (Fig. S2). The DAPC plot showed a similar result to PCA (Fig. 3B): a clear distinction between northern and southern populations (Fig. 3B). The DAPC cluster membership bar plot (Fig. 3C) shows the probabilities of individual assignment to the different clusters, *e. g.* the individual B5 had almost 50% probability of assignment to geographic population B or C. All populations are separated by distinct genetic clusters except for B and C, which can be admixed.

**Fig. 3.**
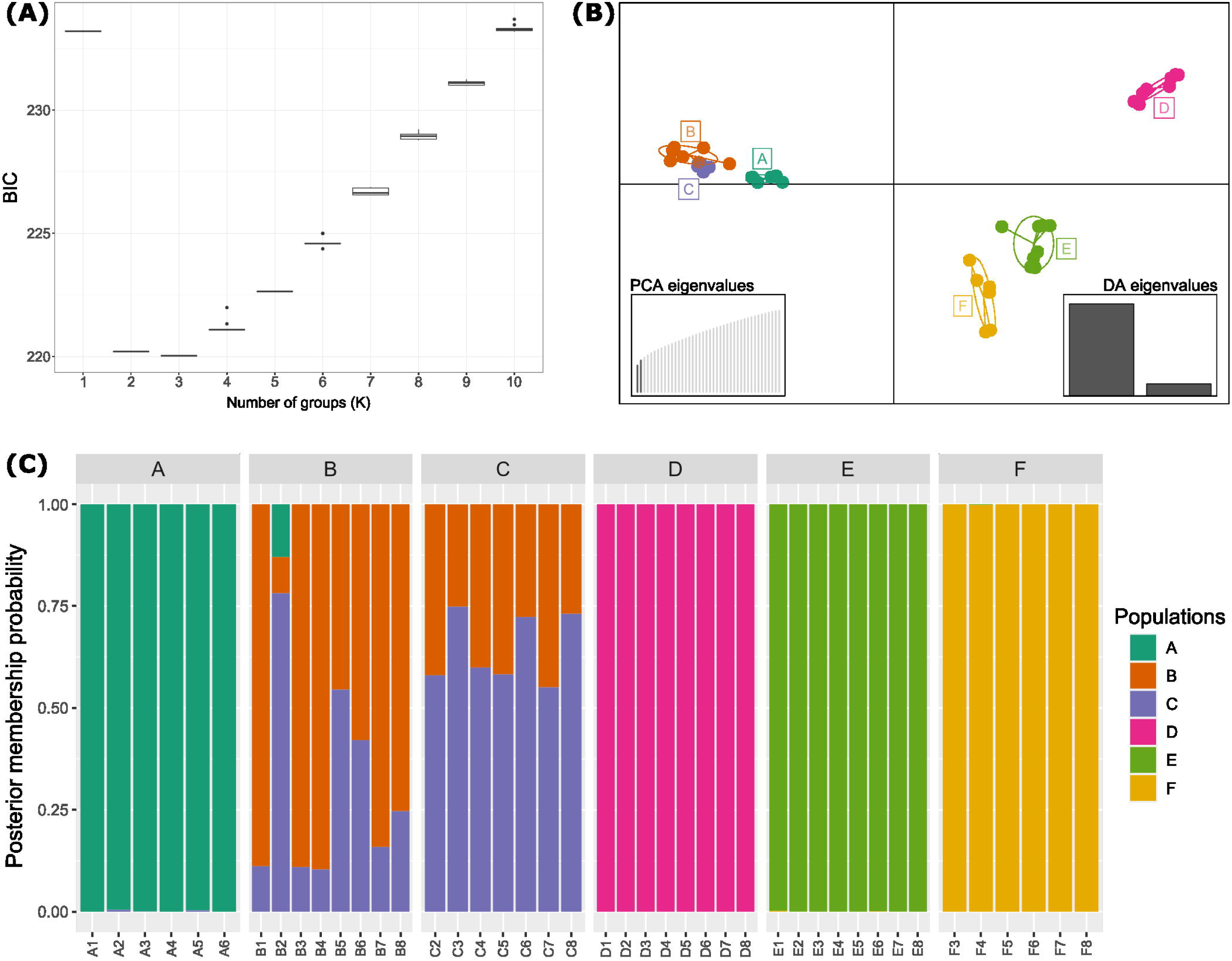
Discriminant Analysis of Principal Components of the genotyping dataset. (A) Bayesian Information Criterion (BIC) values on 1 to 10 connected genetic groups (K). (B) DAPC analysis with retrieving two principal components eigenvalues and two discriminant analysis eigenvalues. Cumulative PCA eigenvalues are shown in the bottom-left graph, in which the first two eigenvalues (dark grey) represent ~39% of the variance. The F-value (2086.6085, 196.4756) of the two discriminant analyses is shown on the bottom-right graph. (C) Composite bar plot showing the posterior probabilities of individual assignment to different clusters, in this case, geographic population. Each individual is indicated per bar (below) and grouped per geographic population site (above). The bar colours indicate the posterior probability proportion of the genetic information from each individual belonging to the genetic population (colour in legend).

### 3.3 Genotypes associated with tolerance to warm temperatures

The *de novo* assembly and genotyping performed with 34 samples, including seven thermally tolerant individuals, resulted in 61,367 SNPs. After filtering, 14,023 SNPs were retained. Our GEMMA analysis indicated that two SNPs are significantly associated with tolerance to warm temperatures (q-value < 0.05; Fig. 4). Individuals not tolerant to warm temperatures did not contain the warm tolerate allele for either SNP, i.e., were homozygous for the non-warm tolerant allele (Table 3). For SNP 287166:52, the warm tolerant allele was found only in the southern population warm tolerant individuals (D2, E5 and E6), while for SNP 7024:55, the warm tolerant allele was found only in warm tolerant individuals from the north (B7, C3 and C8). Individual C2 belonged to a strain line that showed to be warm tolerant (Layton & Johnson, 2021) but did not contain the warm tolerant allele for either of the two SNPs identified here, i.e., was homozygous for the non-warm tolerate allele at both SNPs. We checked if these SNPs were localised in a genomic region with functional meaning, *e.g.,* within or close to a coding gene, but relationships to known coding regions were not found.

**Fig. 4.**
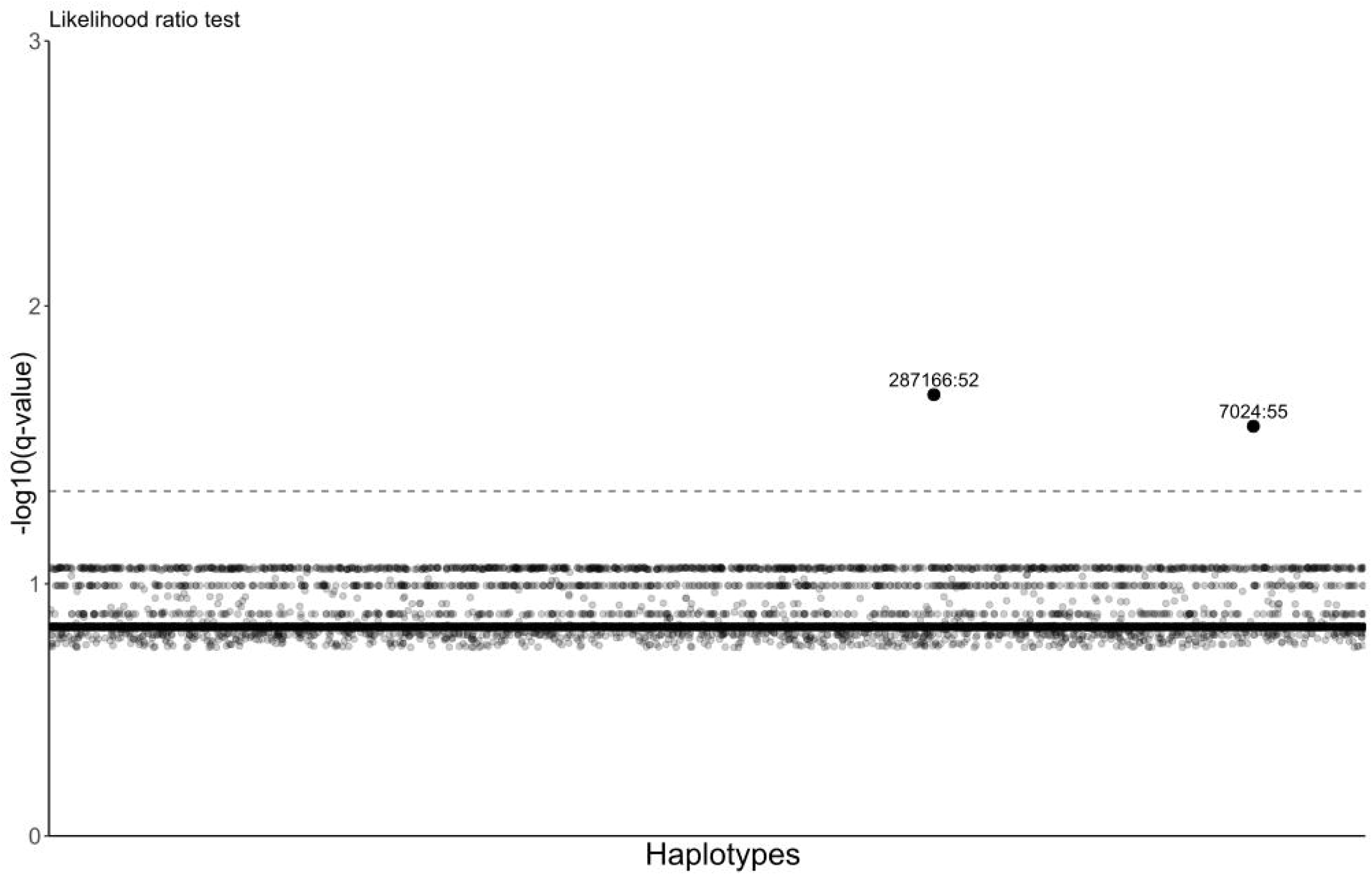
Linear mixed model using likelihood ratio test to identify SNPs significantly associated with warm temperatures. The dashed line indicates the threshold for q-value = 0.05. The two significant SNPs are shown above the threshold.

**Table 3 –.** SNP genotypes for the SNPs associated with tolerance to warm temperatures. 0 = reference (non-warm tolerant) allele. 1 = alternate (warm tolerant) allele. NA = not genotyped.

| Individuals | Genetic group | Phenotype | 287166:52 | 7024:55 |
| --- | --- | --- | --- | --- |
| A1 | North | Non-tolerant | 0/0 | 0/0 |
| B1 | North | Non-tolerant | 0/0 | 0/0 |
| B2 | North | Non-tolerant | NA | NA |
| B3 | North | Non-tolerant | 0/0 | 0/0 |
| B4 | North | Non-tolerant | 0/0 | 0/0 |
| B5 | North | Non-tolerant | 0/0 | 0/0 |
| B6 | North | Non-tolerant | 0/0 | 0/0 |
| B7 | North | Tolerant | 0/0 | **0/1** |
| B8 | North | Non-tolerant | 0/0 | 0/0 |
| C2 | North | Tolerant | 0/0 | **0/0** |
| C3 | North | Tolerant | 0/0 | **0/1** |
| C5 | North | Non-tolerant | 0/0 | 0/0 |
| C6 | North | Non-tolerant | 0/0 | 0/0 |
| C7 | North | Non-tolerant | 0/0 | 0/0 |
| C8 | North | Tolerant | 0/0 | **0/1** |
| D1 | South | Non-tolerant | 0/0 | 0/0 |
| D2 | South | Tolerant | **0/1** | 0/0 |
| D3 | South | Non-tolerant | 0/0 | 0/0 |
| D4 | South | Non-tolerant | 0/0 | 0/0 |
| D5 | South | Non-tolerant | 0/0 | 0/0 |
| D7 | South | Non-tolerant | 0/0 | 0/0 |
| D8 | South | Non-tolerant | 0/0 | 0/0 |
| E1 | South | Non-tolerant | 0/0 | 0/0 |
| E2 | South | Non-tolerant | 0/0 | 0/0 |
| E4 | South | Non-tolerant | 0/0 | 0/0 |
| E5 | South | Tolerant | **0/1** | 0/0 |
| E6 | South | Tolerant | **0/1** | 0/0 |
| E7 | South | Non-tolerant | 0/0 | 0/0 |
| E8 | South | Non-tolerant | 0/0 | 0/0 |
| F3 | South | Non-tolerant | 0/0 | 0/0 |
| F4 | South | Non-tolerant | 0/0 | 0/0 |
| F5 | South | Non-tolerant | 0/0 | 0/0 |
| F6 | South | Non-tolerant | 0/0 | 0/0 |
| F7 | South | Non-tolerant | 0/0 | 0/0 |

## 4 Discussion

### 4.1 Genetic structure of Tasmanian *Macrocystis pyrifera*

Our genetic analysis indicated a strong genetic structure in the *M. pyrifera* populations in Tasmania, with lower connectivity between north and south regions than between populations within either region. This is indicated by the genetic variation explained mostly between regions (Table 2) and the higher F_ST_ between populations from different regions (Fig. 2). Our DAPC analysis also reinforces a genetic distinction between northern versus southern regions in Tasmania (Fig. 3, Fig. S1). Our findings differ from prior genetic analysis using only conservative mitochondrial and plastid molecular markers that revealed negligible genetic diversity among the population in Tasmania (Macaya & Zuccarello, 2010, Durrant *et al*., 2015). A genome-wide approach is more powerful in revealing fine-scale intraspecific structures (Parvizi *et al*., 2022). Distinct genetic structuring using nuclear microsatellite loci (Alberto *et al*., 2009) has, however, been observed in population genetic analyses of *M. pyrifera* in the northeastern and southeastern Pacific (Alberto *et al*., 2010, Johansson *et al*., 2015, Camus *et al*., 2018). These studies showed high genetic differentiation among populations, often over relatively small spatial scales, coinciding with their geographic distribution and a pattern of isolation by distance (Alberto *et al*., 2010, Camus *et al*., 2018). However, other environmental conditions could explain some genetic differentiation (Johansson *et al*., 2015). Low or uni-directional gene flow among populations could decrease fitness through the introduction of genotypes and phenotypes maladapted to the current or future environmental conditions (Johansson *et al*., 2013).

The observed higher genetic differentiation could be caused by the limited effective dispersal distance of their spores and their brief survival (Reed *et al*., 1991, Reed *et al*., 2006, Layton *et al*., 2022). The ability of *M. pyrifera* to raft over considerable distances for longer periods indicates a strong dispersal potential for adults (Hobday, 2000, Macaya *et al*., 2005, Avila *et al*., 2020, Layton *et al*., 2022). However, rafting of adults among dense, established populations may be unlikely to result in significant gene flow (Schiel & Foster, 2015), because in and around established populations, gametes of locals are most numerous and most likely to ‘win’ new space, dramatically outnumbering and swamping opportunities for the gametes of any rafting adult immigrants to the area (see Waters et al. 2013, Fraser et al. 2017). Instead, rafting provides a powerful means of colonisation of new, vacant territory when opportunity allows (Fraser *et al*., 2017, Peters *et al*., 2020). Dispersing adult kelp should be able to seed the colonisation of new space freed by disturbance (Peters *et al*. 2020), such as population crashes in the wake of disturbances. In Tasmania, the more northern populations are more likely to be disturbed by warming impacts (Johnson *et al*., 2011, Butler *et al*., 2020). As surface water flow trends along the Tasmanian eastern coast are mostly southward (Oliver & Holbrook, 2018), there is, therefore, little opportunity for kelp rafting from the denser and generally ‘healthier’ southern populations to replenish the more impacted northern populations. Anthropogenic translocations from south to north could help to counteract this shortcoming. Perhaps sampling a central region (*i.e.,* less distance between regions) would have revealed a transition zone between the severely degraded and fragmented northern region and the more intact and stable areas in the southern region. However, this putative central region has almost no giant kelp remaining. Hence initial sampling did not focus on that area. We note that the southern region had higher differentiation between populations relative to the northern region (Fig. 3), although population-level genetic diversity was similar within the region (Table 1). This higher structure in the south is perhaps evident of historical stability and of a greater number of remnant populations that are surviving in this cooler region, compared to northern populations, which have experienced a longer history of temperature stress, declines, and thus potential bottlenecks (Johnson *et al*., 2011, Butler *et al*., 2020). Such gradients in genetic structure have been noted for other taxa in long-established versus younger populations, for example, with higher differentiation in genetic structure in long-term ‘refugial’ populations than in those that recently recolonised more polar regions in the wake of past ice ages (Hewitt, 2000, Fraser *et al*., 2012).

### 4.2 Scanning genotypes associated with tolerance to warm temperatures

Our genotype-phenotype association results indicate that two loci correlate with warm temperature tolerance (Fig. 4) and thus might be markers that could enable the selection of warm-tolerant lineages for conservation and reforestation efforts. The loci for both SNPs associated with tolerance to warm temperatures were heterozygous in the thermally tolerant individuals. With our limited data set, we could not find robust links between these SNPs and known coding genes to predict the metabolism behind these loci. However, from a conservation perspective, these loci will provide valuable information for the next reforestation activities in Tasmania. Intriguingly, each genetic group possessed a different putative SNP associated with thermal tolerance (Table 3), which could indicate different pathways to local adaptation to temperature tolerance. Such differences are perhaps also due to the limited gene flow between populations from the north and south of Tasmania. Indeed, temperature stress and limited gene flow have driven local adaptation in other seaweeds (King *et al.,* 2019, Miller *et al.,* 2020).

Further experimental studies could employ more powerful genomic approaches, such as whole genome sequencing and transcriptomic analysis, to identify other loci involved in thermal tolerance as well as enable more detailed inferences of the putatively adaptive roles of these SNPs. Future research conducting phenotypic and genomic analysis of offspring from crosses of temperature-tolerant gametophyte strain lines within and between populations could indicate whether these SNPs were truly associated with temperature tolerance. There is also an opportunity to investigate whether crossing thermally tolerant strain lines from north and south would improve the temperature tolerance in offspring. Our results, nonetheless, highlight how genomic tools can assist management programs in identifying adaptively important loci and estimating neutral population genetic parameters (Allendorf *et al.,* 2010). These preliminary insights into the genomic identification of possible warm-tolerant strains will facilitate future research and support the kelp reforestation program in Tasmania.

## 5 Conclusion

This study addressed important knowledge gaps around population genetic structure and warm tolerant genotypes in giant kelp. Our findings support our hypotheses, revealing a stronger genetic structure among *M. pyrifera* populations in Tasmania than previous studies and providing an initial evaluation for a genetic basis for warm tolerance. These results are an essential first step in understanding the genetic structure amongst the remnant giant kelp populations in Tasmania. Understanding the fine-scale genetic structure of *M. pyrifera* population is crucial for successful kelp reforestation. Further studies are necessary to better understand the genetic association with potential temperature-tolerant individuals and populations. These findings have critical relevance for conservation and future management plans for coastal ecosystems, including selective breeding lineages in Tasmania and all regions where giant kelp forests occur for reforestation.

## 7 Author Contributions

CI, CL, CJ, CIF, AW contributed to the conception and design of the study. CI, CL, AW participated in the data collection, sample processing, and conducted the experiments. CI, CAE analysed the data. CI, CL, CIF, CJ, AW contributed to the interpretation of the results and wrote the first draft of the manuscript. All authors contributed to revising and approving the submitted version of the manuscript.

## Acknowledgments

CI and AW would like to thank Dr William Pearman and Prof Melinda Coleman for suggestions for DNA extraction and genotyping sequencing. We gratefully thank Dr Rebecca Jordan for suggestions for the genotype-phenotype association analysis and for reviewing and providing excellent comments for the manuscript. CI would like to thank Neil Holbrook for helpful information about Tasmanian sea currents. Sequencing was performed at the Australian Genome Research Facility.

## 10 Contribution to the field statement

Tasmanian coasts used to feature dense giant kelp forests that severely declined over the last half-century due to changes in environmental conditions, such as ocean warming. Efforts are being made to restore these kelp forests, using strain lines from remnant populations that are potentially tolerant to warm temperatures. However, little is known about the genetic connectivity among these populations, which is essential for conserving the existing forests and future reforestation planning. Here, we present the genetic structure of remnant giant kelp populations from the northern and southern regions of Tasmania and provide an initial genetic basis for the potentially warm-tolerant strains. Using an efficient genotyping approach, we found that the northern and southern populations are distinct genetic groups. Two loci in the genome are significantly associated with tolerance to warm temperatures, although this result is faint, and more research should be conducted to confirm this association. This research is the first step in understanding the genetic diversity amongst giant kelp populations in Tasmania and is critical for conservation and management plans for giant kelp forests.

## 11 Data Availability Statement

The raw data of the *Macrocystis pyrifera* genome and genotyping is available at the European Nucleotide Archive (ENA) with the project accession number ENA: PRJEB55054. Whole genome sequencing raw data is available with the accession numbers ENA: ERR10320122. Genotyping-by-sequencing raw data is available with the accession numbers ENA: ERR10142376-ERR10142419. The genomic and genotyping datasets generated for this study can be found in figshare: https://figshare.com/s/10d87c46afe8d98f8eea (*This is a temporary link for the review process. A public DOI will be generated with the acceptance of the manuscript.)*

