## Supplementary figures and images for "Population genomic analysis reveals genetic structure and thermal-tolerant genotypes in remnant Tasmanian giant kelp populations"

### Fig. S1

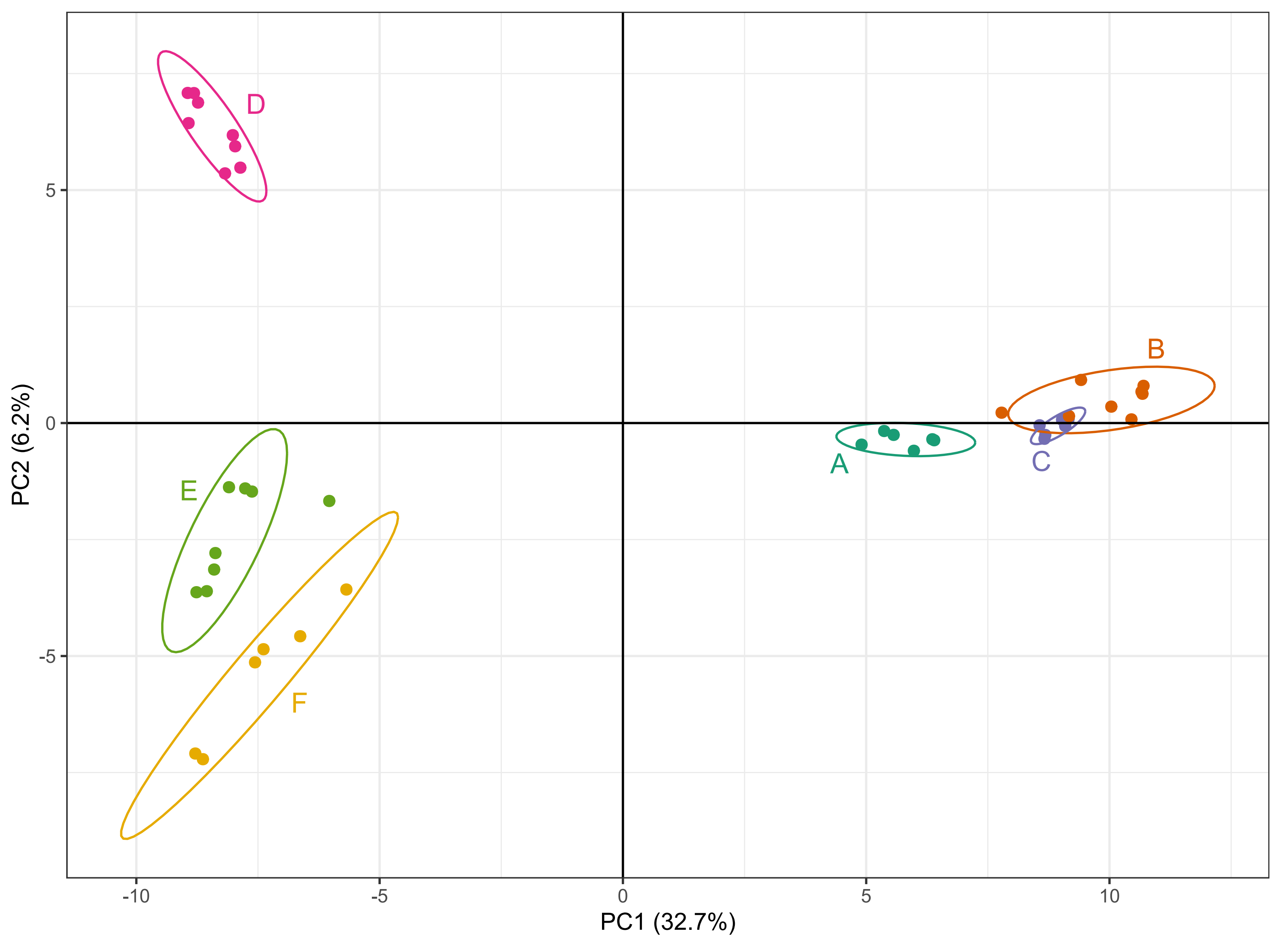

### Fig. S2

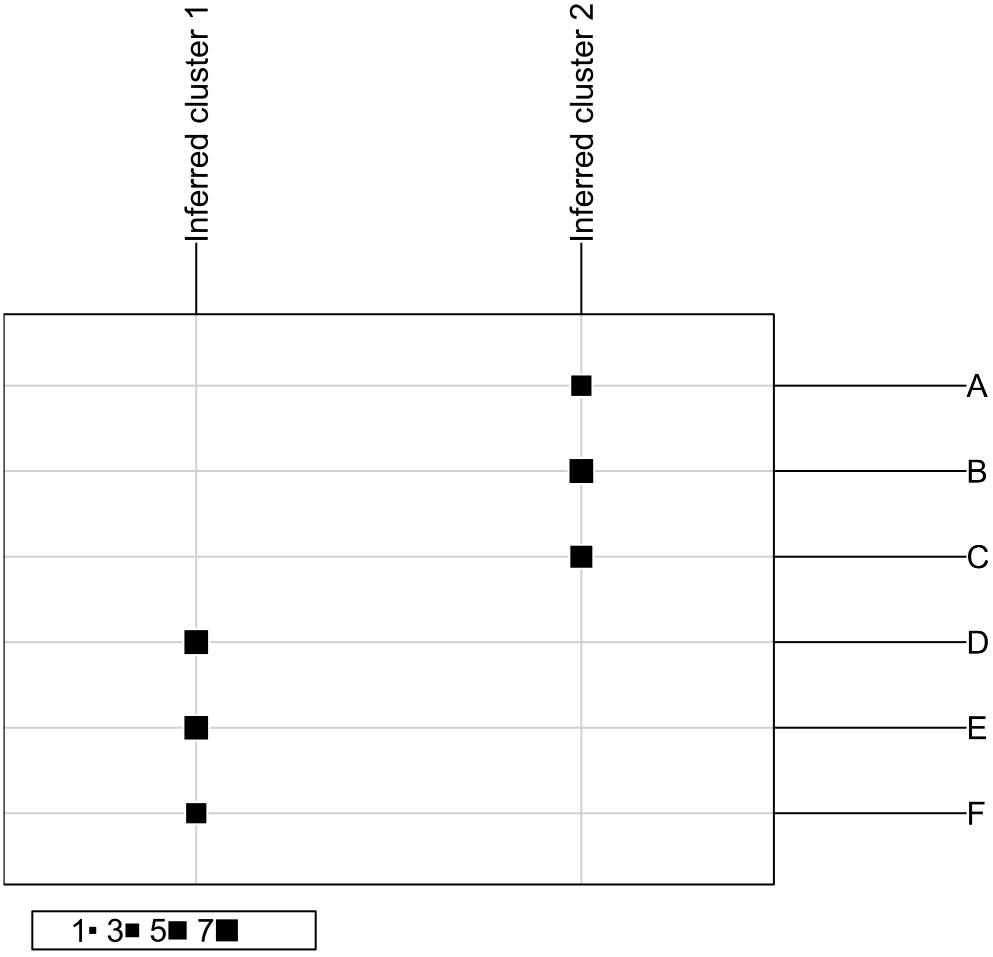
